# In vitro evaluation of antagonism by *Trichoderma* spp. towards *Phellinus noxius* associated with rain tree (*Samanea saman*) and Senegal mahogany (*Khaya senegalensis*) in Singapore

**DOI:** 10.1101/151753

**Authors:** Daniel C. Burcham, Jia Yih Wong, Nelson V. Abarrientos, Mohamed Ismail Mohamed Ali, Yok King Fong, Francis W. M. R. Schwarze

## Abstract

A series of laboratory tests was conducted to screen and identify *Trichoderma* spp. for the biological control of *P. noxius* on pruning wounds. In total, 7 *Trichoderma* isolates were evaluated for their biological fitness and antagonism towards 2 different *P. noxius* isolates associated with Senegal mahogany and rain tree in Singapore. The competitive ability of various *Trichoderma* isolates was assessed by their germination rates, growth rates, and chlamydospore production; and the antagonism of *P. noxius* by *Trichoderma* was assessed by the interaction of these fungi in dual culture and on wood blocks. In this study, the *Trichoderma* isolates showed greater competitiveness and antagonism towards *P. noxius* compared to similar reports in the literature. A majority of the *Trichoderma* isolates germinated and grew at consistently high rates, but there was considerable variability in the production of chlamydospores among isolates. All *Trichoderma* isolates routinely antagonized *P. noxius* in the different bioassays, but there was significant variability in the antagonistic capacity of various isolates and in the susceptibility of the *P. noxius* isolates to confrontation with *Trichoderma.* Relative to the controls, *P. noxius* cultures grew significantly slower in the presence of volatile organic compounds emitted by most *Trichoderma* isolates, and a large majority of *Trichoderma* isolates caused a significant reduction to the dry weight loss of wood blocks inoculated with *P. noxius.* Although no single *Trichoderma* isolate consistently antagonized *P. noxius* better than all others in every test, *T. harzianum* 9132 and *T. virens* W23 notably did so more regularly than others.

## Introduction

Trees are pruned for many reasons, including aesthetics, clearance, and safety. However, the exposed wood on pruning wounds is susceptible to wood decay infection (Barry et al., 2000; Deflorio et al., 2007), and these infections are a concern to people interested in preserving the environmental benefits and commercial value of trees. Historically, arborists often applied various treatments to pruning wounds to create a physical barrier or inhibit wood decay fungi (Lonsdale, 1984). Although some of these treatments improved wound occlusion (Mercer, 1983) and temporarily prevented infection (Mercer et al., 1983), the physical barriers frequently deteriorated over time due to weathering, insect damage, and growth stress that, in some cases, exacerbated decay compared to untreated wounds (Mercer et al., 1983). Similarly, some fungicides were reported as effective at limiting infection but their phytotoxicity often caused harmful cambial dieback (Mercer et al., 1983; Mercer, 1979) that inhibited wound occlusion and increased wood decay by expanding the exposed wound surface. In addition to their limited efficacy, the use of synthetic chemicals, especially on trees planted in densely populated urban areas, creates environmental and human health risks that many consider unacceptable. As a result, most arboriculture industry standards discourage wound treatment (Anonymous 2008).

Biological control agents, however, offer a promising alternative to traditional wound treatments. In particular, many fungal antagonists have been selected from the genus *Trichoderma* for biological control of plant diseases, including economically important fungal pathogens affecting fruit trees (Ricard and Highley, 1988) and conifers (Kallio and Hallaksela, 1979). Notably, Pottle and Shigo (1975) reported that basidiomycetes were absent from 1-year-old red maple *[Acer rubrum* L. (Sapindaceae)] wounds preventatively treated with *T. viride,* but decay fungi were present in 80% of untreated control wounds after the same period. Mercer and Kirk (1984b) similarly reported a significant reduction in wood decay fungi infection of European beech *[Fagus sylvatica* L. (Fagaceae)] wounds treated with a different isolate of *T. viride,* compared to control wounds. Notably, *T. viride* was more effective and persisted longer than other biological controls (Mercer and Kirk, 1984b). More recently, Schubert et al. (2008a) demonstrated the ability of a *T. atroviride* isolate to significantly reduce wood decay infection rates among more than 1,400 wounds treated from 6 different broadleaf tree species.

*Trichoderma* spp. (Samuels, 1996) are ubiquitous, globally distributed saprophytes that occur primarily in highly organic soils (Papavizas, 1985), but they are capable of colonizing a wide range of natural substrates (Klein and Eveleigh, 1998). Many *Trichoderma* spp. are avirulent plant symbionts (Harman et al., 2004) that antagonize phytopathogenic fungi through mycoparasitism (Lorito et al., 1996a), antibiosis (Ghisalberti and Sivasithamparam, 1991), enzyme production (Markovich and Kononova, 2003), and competition for resources (Sivan and Chet, 1989). In most cases, a single *Trichoderma* spp. isolate simultaneously employs more than one of these antagonistic mechanisms to suppress disease-causing fungi; there are several reports of synergistic enhancement between multiple antagonistic processes and substances (Di Pietro et al., 1993; Lorito et al., 1994a; Lorito et al., 1994b; Lorito et al., 1996b). In addition to suppressing harmful microbes, soil application of *Trichoderma* spp. enhances root growth and nutrient uptake efficiency, resulting in increased overall plant growth and reproductive yield (Harman et al., 2004). Several authors have also demonstrated that some *Trichoderma* spp. cause systemic induced resistance (Pieterse and van Loon, 1999) in plants by observing physical separation between the introduced *Trichoderma* spp. and area of disease reduction (Bigirimana et al., 1997; Djonovic et al., 2006; Yedidia et al., 1999).

Similar to mycorrhizae, many benefits accrue to plants after *Trichoderma* spp. colonize the outermost epidermal layers of their roots (Yedidia et al., 2000). Although most *Trichoderma* spp. are avirulent, a few isolates are pathogenic or phytotoxic (Koike et al., 2001), and it is important to screen for potential biological control agents with representative laboratory tests that mimic the temperature, moisture, and nutrient availability conditions expected for the desired field application (Howell, 2003). Moreover, many authors caution that individual *Trichoderma* spp. isolates usually do not have the intrinsic ability to exert all of the reported biocontrol mechanisms (Harman, 2000; Howell, 2003), and hybridization, genetic modification, or mixing may be needed to combine the desirable characteristics of multiple isolates (Brunner et al., 2005; Djonovic et al., 2007).

However, most studies addressing biological control with *Trichoderma* have been conducted in temperate climates with associated species. *Phellinus noxius* (Corner) G. Cunn (Hymenochaetaceae) is an aggressive, often lethal wood decay fungus that infects the root systems of many tree species throughout the global tropics and subtropics (Rossman and Farr, 2014). Macroscopically, the wood of infected trees appears brown and the adjacent root flare is often covered by a dark brown mycelial mat (Chang, 1995). Although the disease is commonly called brown root rot, *P. noxius* actually causes a simultaneous rot by degrading cellulose and lignin at the same time (Nicole et al., 1995). A few recent studies have evaluated the ability of *Trichoderma* to antagonize *P. noxius* obtained from root system infections in Australia (Schwarze et al., 2012) and Hong Kong (Ribera et al., 2016). Recent work in Singapore, however, demonstrated the ability of *P. noxius* to colonize pruning wounds, invade host defensive responses, and cause severe wood decay in the aboveground parts of Senegal mahogany *[Khaya senegalensis* (Desr.) A. Juss. (Meliaceae)]; this work revealed that *P. noxius* occupies a broader ecological niche than previously understood with physiological capacity for multiple trophic strategies (Burcham et al., 2015). At the same time, diagnostic efforts have shown that *P. noxius* similarly infects other trees in Singapore, including rain tree *[Samanea saman* (Jacq.) Merr. (Fabaceae)].

Collectively, Senegal mahogany and rain tree comprise more than 10% of Singapore’s urban forest and generate a significant part of the total benefits rendered to the surrounding community. In general, the ubiquity of pruning wounds creates considerable opportunity for *P. noxius* infections, and limiting these infections is an important strategy to protect susceptible trees and, by extension, the health of the urban forest. As a result, the objectives of this study were (1) to screen *Trichoderma* spp. isolates for antagonistic potential against *P. noxius* associated with pruning wounds on Senegal mahogany and rain tree and (2) to identify the most competitive isolate(s) for field testing. To this end, a series of bioassays were designed to evaluate antagonism by *Trichoderma* spp. isolates towards *P. noxius* in laboratory conditions that resembled situations expected for landscape application in Singapore.

## Material and methods

### Origin of fungal isolates

*P. noxius* isolates were obtained from decayed wood columns that were excised near existing branch pruning wounds on Senegal mahogany and rain tree in Singapore. Among several obtained from these tree species, the most aggressive *P. noxius* isolate was selected for screening biological control agents based on its superior ability to cause dry weight loss to species-specific sapwood blocks in laboratory tests (data not shown). In total, two *P. noxius* isolates were selected–one caused maximum dry weight loss to Senegal mahogany and the other to rain tree wood blocks. Separately, 7 *Trichoderma* spp. isolates were obtained from the fruiting bodies of basidiomycetes colonizing various plant species and organic substrates in Singapore. The identity and origin of fungal isolates used in this study is given in Table 1.

**Table 1.** Identity, origin, and accession numbers of the fungal isolates collected for use in this study.

| Species | Isolate | Accession No. | Substrate | Origin | Reference | Identity (%) |
| --- | --- | --- | --- | --- | --- | --- |
| **a. Wood decay fungi** |  |  |  |  |  |  |
| *P. noxius* | KSK6 | KF233592 | ex. *Khaya senegalensis* | Bugis,<br>Singapore | FR821770<br>(Schwarze et al., 2012) | 99 |
| *P. noxius* | SSN3P | KP202302 | ex. *Samanea saman* | Clementi,<br>Singapore | FR821769<br>(Schwarze et al., 2012) | 98 |
| **b. *Trichoderma* spp.** |  |  |  |  |  |  |
| *T. reesei* | 1056TD1 | KY025557 | *Ganoderma boninense*<br>on *Cocos nucifera* | Singapore | JQ979435.1<br>(Kuhls et al., 1996) | 99 |
| *T. asperellum* | 3 | KY025555 | *Ganoderma* sp.<br>on coarse woody debris | Bukit Batok,<br>Singapore | AJ230669.1<br>(Lieckfeldt et al., 1999) | 100 |
| *T. harzianum* | 9132 | KY025556 | *Ganoderma boninense*<br>on *Cyrtostachys renda* | Singapore | KR856210.1<br>(Lieckfeldt et al., 2001) | 99 |
| *T. saturnisporum* | SPH20109132 | KY025558 | *Ganoderma boninense*<br>on *Cyrtostachys renda* | Singapore | NR_103704.1<br>(Kuhls et al., 1997) | 99 |
| *T. harzianum* | W21 | KY025559 | *Ganoderma boninense*<br>on *Ptychosperma macarthurii* | Woodlands,<br>Singapore | KJ000324.1<br>(Lieckfeldt et al., 2001) | 99 |
| *T. virens* | W23 | KY025560 | *Ganoderma boninense*<br>on *Ptychosperma macarthurii* | Woodlands,<br>Singapore | KP009289.1<br>(Ottenheim et al., 2015) | 100 |
| *T. virens* | W26 | KY025561 | *Ganoderma boninense*<br>on *Ptychosperma macarthurii* | Woodlands,<br>Singapore | KR296867.1<br>(Ottenheim et al., 2015) | 100 |

Note: Fungal ITS sequences were deposited in GenBank (Benson et al., 1997).

### Cultural media and growth conditions

Three cultural media were used, including a malt extract agar (MEA, Thermo Fisher Scientific, Waltham, MA, USA); a low nutrient agar (LNA) prepared according to Huttermann and Volger (1973), with each liter containing 0.013 g H2O: L-asparagine, 1 g KH2PO4, 0.3 g MgSO4, 0.5 g KCL, 0.01 g FeSO_4_, 0.008 g MnSO_4_ 4H_2_O, 0.002 g ZnSO_4_ 6H_2_O, 0.05 g CaNO_3_ 4H2O, 0.002 g CuSO_4_, 0.008 g NH4NO3, 5 g D-glucose, and 10 g agar; and a basidiomycete selective medium (BSM) modified from Sieber (1995), with each liter of media containing 50 g MEA, 105.75 mg thiabendazole dissolved in 2 ml of concentrated lactic acid, 200 mg chloramphenicol, and 300 mg streptomycin sulfate. Unless indicated otherwise, cultures were incubated consistently in the dark at 28 °C and 50–70% RH on 90 or 50 mm Petri dishes containing approximately 20 or 11 ml, respectively, of media. All plates were inoculated with 8 mm mycelial discs extracted from the actively growing margins of pure cultures on MEA and sealed with plastic paraffin film (Parafilm®, Pechiney Plastic Packaging, Chicago, IL, USA).

### Germination and growth rates

Growth rates of the *Trichoderma* isolates were evaluated on MEA and LNA. The LNA was chosen because its higher C:N ratio better replicated the nutritional status of wood (Srinivasan et al., 1992) than MEA. To assess growth, 8 mm mycelial discs of *Trichoderma* were inoculated centrally onto MEA or LNA in a 90 mm Petri dish. Growth rates were determined by recording the average of 2 orthogonal fungal culture diameter measurements at regular intervals over a 36-hour period.

To determine germination rates, a conidial suspension of each *Trichoderma* isolate was prepared by flooding mature cultures with sterile water, dislodging conidia by physical agitation, and skimming buoyant conidia from the surface. The concentration of suspensions was assessed using a haemocytometer and adjusted to obtain approximately 10^5^ colony forming units (CFU) ·ml^-1^. Subsequently, 10 μl of each suspension was placed in 3 separate locations on a single Petri dish containing LNA. Test plates were covered, sealed with paraffin tape, and incubated at constant conditions for 18 hours. After incubation, conidia were observed at 10× magnification with a light microscope (Nikon Eclipse E200, Tokyo, Japan) in 5 randomly selected locations in each Petri dish. The germination rate was calculated for each plate as the average percent of germinated conidia contained in these fields of observation.

### Chlamydospore production

To assess the production of thick-walled resting spores during adverse environmental conditions, chlamydospores were counted on 14-day old mature cultures of *Trichoderma* growing on LNA in a 50 mm Petri dish. The number of chlamydospores was determined by counting those observed at 10× magnification in 5 randomly selected locations on each Petri dish using a digital camera system (Nikon Digital Sight DS-Fi1, Tokyo, Japan) attached to a light microscope. The number of chlamydospores on each plate was computed as the average number observed across the different locations.

### VOC inhibition

Inhibition of wood decay fungi by volatile organic compounds (VOCs) produced by *Trichoderma* isolates was evaluated using tests described by Dennis and Webster (1971). Initially, 8 mm mycelial discs of *Trichoderma* isolates were placed in the center of a Petri dish containing MEA. At the same time, each of the *P. noxius* isolates was inoculated identically onto a separate Petri dish. Each *Trichoderma* isolate was then paired with a *P. noxius* isolate by inverting the bottom of one Petri dish containing *Trichoderma* and replacing the top of another containing *P. noxius.* The rims of the paired Petri dishes were aligned and sealed. The diameter of *P. noxius* cultures was recorded as the average of 2 orthogonal measurements at regular 24-hour intervals over a 5-day period. Similar Petri dishes without *Trichoderma* isolates served as a control.

### Interaction tests

Mycoparasitism was assessed in dual culture and on wood blocks broadly according to Highley et al. (1997) and Schubert et al. (2008a), respectively. For dual culture tests, 8 mm mycelial discs were removed from 14-day old, actively growing pure cultures of each *P. noxius* isolate and placed individually at a regular distance (~ 10 mm) from the edge of a 90 mm Petri dish containing MEA. After incubation for 72 hours, equivalent mycelial discs were removed from 7-day old actively growing pure cultures of each *Trichoderma* isolate, inserted on a Petri dish opposite the growing wood decay fungus, covered, and sealed. The Petri dishes were incubated under constant conditions for 4 weeks. The ability of *Trichoderma* isolates to parasitize *P. noxius* isolates was assessed by removing 6 mycelial discs from the final extent of the wood day fungus culture and placing discs on BSM. The media’s inhibition of *Trichoderma* allowed a selective determination of the lethal effect of *Trichoderma* isolates, determined as one minus the percent of these 6 discs yielding *P. noxius.*

For wood block interaction tests, wood blocks (10 × 8 × 30 mm) were removed from healthy sapwood in Senegal mahogany and rain tree. Individual wood blocks were oven-dried at 100 °C for 48 hours, cooled in a vacuum desiccator, and weighed using a precision balance. The blocks were subsequently autoclaved twice at 121 °C for 30 min. Each wood block was inoculated by immersion for 10 minutes in a conidial suspension containing 10^5^ CFU ml^-1^ of a *Trichoderma* isolate, 0.2% D-glucose, and 0.1% urea. The inoculated wood blocks were incubated at constant conditions in a 50 ml sterile centrifuge tube for 4 weeks. Afterwards, the Senegal mahogany and rain tree wood blocks pre-treated with *Trichoderma* were placed on a 14-day old pure culture of one *P. noxius* isolate growing on MEA in a 50 mm Petri dish. Test plates were covered, sealed, and incubated under constant conditions for 12 weeks. Untreated, sterilized wood blocks were similarly placed on 14-day old actively growing pure cultures of *P. noxius* and served as controls. After incubation, the wood blocks were cleaned by removing surface mycelia, oven-dried, cooled, and reweighed. The dry-weight loss was then determined for each wood block on a percentage basis.

### Statistical analysis

For all experiments, 5 replicates were maintained of each treatment combination. Since germination rates and chlamydospore counts were not measured under experimentally manipulated conditions, the results were not statistically analyzed but instead presented as descriptive data. The effect of experimental treatments on the measured response was determined using analysis of variance in SAS 9.4 (SAS Institute, Inc., Cary, NC, USA). For growth rate bioassays, model fixed effects included media type, *Trichoderma* isolate, and their interaction. Since the rate of change in culture diameter with respect to time was constant, growth rates were analyzed as the slope of a linear function (Gams and Bissett, 1998) fit to paired observations of culture diameter and time using least squares regression. For the remaining bioassays, model fixed effects included *Trichoderma* isolates, *P. noxius* isolates, and their interaction. Given our primary interest in antagonism by *Trichoderma* isolates, significant interactions were separated to determine differences among *Trichoderma* isolates within specific levels of the other fixed effect. Mean separation was performed using Dunnett’s one-way test and Tukey’s honestly significant difference for experiments with and without controls, respectively.

## Results

Except for *T. reesei* 1056TD1, all *Trichoderma* isolates germinated, on average, at nearly maximum rates between 99 and 100%. Still, light microscopy revealed that a considerable majority (87%) of *T. reesei* 1056TD1 conidia germinated on LNA. In contrast, there was sizeable variability in the production of chlamydospores among *Trichoderma* isolates. The average number of chlamydospores observed among 5 randomly-selected visual fields at 10× magnification on each Petri dish ranged between 6 (SD 2) for *T. saturnisporum* SPH20109132 and 189 (SD 30) for *T. virens* W23 (Table 2).

**Table 2.** Germination rates (%) and chlamydospore counts (n) for various *Trichoderma* spp. isolates cultured on a low nutrient agar (LNA)

| *Trichoderma* spp. | Germination rate (%) | Chlamydospore count (*n*) |
| --- | --- | --- |
| 1056TD1 | 87 (21) | 9 (4) |
| 3 | 100 (0) | 20 (15) |
| 9132 | 100 (0) | 20 (13) |
| SPH20109132 | 100 (0) | 6 (2) |
| W21 | 100 (0) | 39 (8) |
| W23 | 99 (1) | 189 (30) |
| W26 | 99 (1) | 165 (44) |

Note: Values listed are mean (SD). See Table 1 for the taxonomic identity of fungal isolates.

Overall, there were significant differences in growth rates among *Trichoderma* isolates *(F* = 103.46; df = 6, 24; *p* < 0.001), and these growth rates differed significantly between the 2 media types (F= 107.83; df = 1, 24; *p* < 0.001). Specifically, *Trichoderma* isolates grew significantly faster on MEA than on LNA (Table 3). However, *Trichoderma* isolates and media types interacted significantly to affect growth rates *(F* = 34.55; df = 6, 24; *p* < 0.001). There were significant differences in growth rates among *Trichoderma* isolates growing on both LNA and MEA, but the relative ranking of *Trichoderma* isolates within each media type was not consistent (Table 3). Notably, the growth rate of *T. virens* W23 was significantly slower than all other *Trichoderma* isolates, except *T. virens* W26, on MEA, but its growth rate actually increased by 8.0% on LNA to a value not significantly different from the 4 other fastest growing isolates. The growth rate of *T. harzianum* 9132 decreased marginally by only 2.9% on LNA compared to MEA; the other *Trichoderma* isolates grew between 11.6 and 40.2% slower on LNA (Table 3).

**Table 3.** Growth rates (cm-day^-1^) of *Trichoderma* spp. isolates cultured on 2 different media, including a malt extract agar (MEA) and low nutrient agar (LNA)

| *Trichoderma* spp. | Media type |  |
| --- | --- | --- |
|  | MEA | LNA |
| 1056TD1 | 7.96 (0.11) a | 4.76 (0.04) a |
| 3 | 4.61 (0.01) c | 3.12 (0.27) c |
| 9132 | 4.74 (0.02) c | 4.60 (0.02) ab |
| SPH20109132 | 8.04 (0.10) a | 5.28 (0.48) a |
| W21 | 5.07 (0.11) b | 4.48 (0.76) ab |
| W23 | 4.38 (0.10) d | 4.73 (0.07) a |
| W26 | 4.41 (0.08) d | 3.62 (0.97) bc |
| Mean | 5.61 (1.56) | 4.37 (0.84) |

Note: Values listed are mean (SD). Means followed by the same letter within each column are not significantly different at the $\alpha = 0.05$ level. See Table 1 for the taxonomic identity of fungal isolates.

Compared to the controls, all *P. noxius* cultures grew slower, in absolute terms, in the presence of VOCs emitted by *Trichoderma.* Overall, growth rates of *P. noxius* varied significantly in the presence of VOCs produced by *Trichoderma* isolates (F = 5.40; df = 7, 28; *p* < 0.001), and the growth of *P. noxius* SSN3P cultures was inhibited significantly more by *Trichoderma* VOC emission than *P. noxius* KSK6 (F = 24.14; df = 1, 28; *P* = 0.008). However, *Trichoderma* and *P. noxius* isolates interacted significantly to affect growth rates because VOCs emitted by *Trichoderma* isolates did not inhibit the 2 *P. noxius* isolates in a proportionally consistent way; *Trichoderma* isolates significantly inhibiting one *P. noxius* isolate did not necessarily inhibit the other equally well. Compared to the controls, 6 *Trichoderma* isolates caused a significant reduction to the growth rate of *P. noxius* SSN3P, but only 4 *Trichoderma* isolates caused similar reductions to *P. noxius* KSK6 (Table 4). Separately, *T. harzianum* 9132 and *T. virens* W23 caused the greatest inhibition to *P. noxius* KSK6 and *P. noxius* SSN3P, respectively (Table 4).

**Table 4.** *Phellinusnoxius* KSK6 and *P. noxius* SSN3P growth rates (cm-day^-1^) in the presence of VOCs emitted by various *Trichoderma* spp. isolates

| *Trichoderma* spp. | *P. noxius* |  |
| --- | --- | --- |
|  | KSK6 | SSN3P |
| Control | 1.75 (0.01) | 1.34 (0.08) |
| 1056TD1 | 1.48 (0.19) | 0.42 (0.47)* |
| 3 | 1.46 (0.09) | 0.68 (0.49)* |
| 9132 | 0.55 (0.45)* | 0.69 (0.29)* |
| SPH20109132 | 0.90 (0.32)* | 0.86 (0.21)* |
| W21 | 1.18 (0.36)* | 1.04 (0.25) |
| W23 | 1.52 (0.23) | 0.25 (0.33)* |
| W26 | 0.94 (0.59)* | 0.39 (0.55)* |
| Mean | 1.22 (0.48) | 0.71 (0.48) |

Note: Values listed are mean (SD). Within each column, asterisks (\*) indicate a significant reduction in growth rates compared to the control at the $\alpha = 0.05$ level. See Table 1 for the taxonomic identity of fungal isolates.

In dual culture tests on MEA, most of the *Trichoderma* isolates exhibited comparably high antagonistic potential with percent mortality to *P. noxius* ranging between 80 and 100%. All *Trichoderma* isolates overspread and colonized the *P. noxius* cultures; none were deadlocked in distant confrontation or evaded by the wood decay fungus. Overall, there were no significant differences in the percent mortality of *P. noxius* challenged by various *Trichoderma* isolates in dual culture tests *(F* = 0.98; df = 6, 12; *p* = 0.479), and there were no significant differences in the susceptibility of the 2 *P. noxius* isolates to antagonism by *Trichoderma (F* = 0.61; df = 1, 12; *p* = 0.517). However, *Trichoderma* and *P. noxius* isolates interacted significantly to affect percent mortality (F = 3.25; df = 6, 12; *p* = 0.039). Although the lethal effect of various *Trichoderma* isolates towards *P. noxius* SSN3P in dual culture did not vary significantly, the same was not true for *P. noxius* KSK6. Specifically, the percent mortality of *P. noxius* KSK6 confronted by *T. harzianum* 9132 was significantly less than that caused by the other *Trichoderma* isolates (Table 5).

**Table 5.** Percent mortality (%) to *Phellinus noxius* KSK6 and *P. noxius* SSN3P due to mycoparasitism by various *Trichoderma* spp. isolates in dual culture

| *Trichoderma* spp. | *P. noxius* |  |
| --- | --- | --- |
|  | KSK6 | SSN3P |
| 1056TD1 | 89 (19) a | 89 (10) |
| 3 | 83 (17) a | 94 (10) |
| 9132 | 33 (58) b | 100 (0) |
| SPH20109132 | 94 (10) a | 89 (10) |
| W21 | 78 (10) a | 94 (10) |
| W23 | 89 (10) a | 67 (58) |
| W26 | 100 (0) a | 100 (0) |
| Mean | 81 (29) | 90 (22) |

Note: Values listed are mean (SD). Within left-hand column, means followed by the same letter are not significantly different at the $\alpha = 0.05$ level. See Table 1 for the taxonomic identity of fungal isolates.

Overall, dry weight loss of wood blocks varied significantly according to their pre-treatment with various *Trichoderma* isolates (F = 7.65; df = 7, 28; *p* < 0.001), and pre-treatment with *Trichoderma* isolates gave significantly better protection against decomposition by *P. noxius* SSN3P than *P. noxius* KSK6 (F = 20.52; df = 1, 28; *p* = 0.011). The interaction between *Trichoderma* and *P. noxius* isolates, however, was not significant (F = 1.11; df = 7, 28; *p* = 0.386) because *Trichoderma* isolates inhibited decomposition by the 2 *P. noxius* isolates similarly. Compared to the controls, pre-treatment with 6 different *Trichoderma* isolates caused a significant reduction to the dry weight loss of wood blocks inoculated with *P. noxius.* Among all isolates, *T. harzianum* 9132 offered the greatest protection to wood blocks against decomposition by *P. noxius,* and *T. virens* W26 was the only isolate that did not significantly inhibit wood block decomposition compared to the controls (Table 6).

**Table 6.** Dry weight loss (%) caused to Senegal mahogany *(Khaya senegalensis)* and rain tree *(Samanea saman)* wood blocks by *Phellinus noxius* KSK6 and *P. noxius* SSN3P, respectively, after pre-treatment with various *Trichoderma* spp. isolates

| *Trichoderma* spp. | *P. noxius* |  | Mean |
| --- | --- | --- | --- |
|  | KSK6 | SSN3P |  |
| Control | 12.7 (4.4) | 10.8 (5.0) | 11.7 |
| 1056TD1 | 8.0 (2.6) | 3.6 (5.0) | 5.8* |
| 3 | 4.6 (0.5) | 2.8 (0.8) | 3.7* |
| 9132 | 5.0 (0.8) | 1.8 (1.3) | 3.4* |
| SPH20109132 | 7.7 (2.4) | 4.0 (4.2) | 5.7* |
| W21 | 8.7 (0.8) | 3.6 (2.9) | 6.2* |
| W23 | 7.2 (1.2) | 5.8 (4.2) | 6.5* |
| W26 | 8.2 (2.1) | 8.6 (1.7) | 8.4 |
| Mean | 7.8 (3.1) a | 5.1 (4.3) b |  |

Note: Values listed are mean (SD). Within right-hand column, asterisks (\*) indicate a significant reduction in dry weight loss compared to the control at the $\alpha = 0.05$ level. Means followed by the same letter within the bottom row are not significantly different at the $\alpha = 0.05$ level. See Table 1 for the taxonomic identity of fungal isolates.

## Discussion

Despite apparent variability, all of the *Trichoderma* isolates tested in this study antagonized both *P. noxius* isolates in the different bioassays. In contrast with similar studies (Mercer and Kirk, 1984a; Ribera et al., 2016; Schubert et al., 2008a; Schwarze et al., 2012), none of the *Trichoderma* isolates enhanced the activity of *P. noxius* isolates in culture. Still, no single *Trichoderma* isolate was consistently superior among all others in its ability to antagonize *P. noxius* across every bioassay. In addition, the presence of *Trichoderma* caused a greater reduction to the growth (Table 4) and wood decay rates (Table 6) of *P. noxius* SSN3P than *P. noxius* KSK6, indicating the greater susceptibility of the former to biological control by *Trichoderma.* Although the wood block interaction tests were performed on different wood substrates corresponding to the tree species from which each *P. noxius* isolate was originally obtained (Table 1), the equally greater antagonistic effect of *Trichoderma* VOC emission on *P. noxius* SSN3P suggests that the disparity is not entirely caused by differences in test conditions associated with wood anatomy. In agreement with existing reports (Schwarze et al., 2012; Wells and Bell, 1979), this intraspecific variability and target specificity affirms the importance of screening potential biological control agents against the desired pathogen in laboratory tests.

Germination rates were nearly complete for the conidia of *Trichoderma* isolates cultured on LNA (Table 2). Although exogenous nutrients are required for germination, existing reports indicate that leached nutrients and secondary metabolites can provide adequate resources to support germination (Danielson and Davey, 1973; Naar and Kecskes, 1998). Since conidia were unwashed, germination rates reported in this study may not reflect those for *Trichoderma* in sterile environments. Practically, these results suggest that conidial preparations for biological control can be amended with modest supplemental nutrients to support high germination rates. Analogously, Schubert et al. (2008a) reported that an unmodified conidial suspension and another amended with glucose and urea persisted equally well on tree wounds over a 30-month period. In addition, high germination rates for introduced conidia valuably limits the availability of dead propagules as a nutrient source to other deleterious microbes that might interfere with biological control by competing with *Trichoderma* or infecting the plant host (Harman et al., 1991).

In contrast, however, there was considerable variability in the production of chlamydospores among *Trichoderma* isolates (Table 2). Relatively few reports exist on *Trichoderma* chlamydospore biology; Lewis and Papavizas (1984) reported that their production rates similarly varied among *Trichoderma* isolates. Although it is generally accepted that these resting spores enhance survival during unfavorable conditions in the soil, it is not clear to what extent they might serve a similar purpose on other substrates, such as wood. Practically, chlamydospores are valuable for commercial *Trichoderma* formulations since these propagules better tolerate desiccation and storage than conidia or mycelial fragments (Li et al., 2016).

Growth rates varied significantly among *Trichoderma* isolates, but all isolates grew at faster rates than reported for others selected from temperate or sub-tropical climates for similar biological control applications (Ribera et al., 2016; Schubert et al., 2008a; Schwarze et al., 2012). In agreement with existing work (Schubert et al., 2008b), growth rates decreased for isolates cultured under nutrient limiting conditions on LNA. However, the increased growth of *T. virens* W23 under these conditions is a notable exception (Table 3). Although difficult to demonstrate experimentally (Harman, 2000), competition for space and nutrients is likely an important mechanism for biological control by *Trichoderma*.

It is useful to note that germination and growth rates were determined under constant environmental conditions chosen to replicate typical values for Singapore. At this location, the climate is representative of the equatorial tropics with stable temperatures ranging daily between 24° and 32° C, abundant rainfall approaching 250 cm annually, and elevated humidity between 60 and 90% (Micheline and Ng, 2012). In general, the optimum temperature for the growth of many *Trichoderma* spp. is between 25° and 30° C (Klein and Eveleigh, 1998). There is considerable experimental evidence that *Trichoderma* germination and growth rates are proportional to water activity and temperature (Kredics et al., 2003), up to some limiting value that depends on the climatic origin of a given *Trichoderma* isolate (Ribera et al., 2016; Schubert et al., 2008b; Schwarze et al., 2012); but it was assumed that tolerance to a wide range of unfavorable temperature and moisture conditions is not critical for biological control by *Trichoderma* in Singapore.

Compared to similar work on wood decay fungi (Mercer and Kirk, 1984a; Schubert et al., 2008b), including *P. noxius* (Ribera et al., 2016; Schwarze et al., 2012), VOC emission by *Trichoderma* isolates had a comparatively large effect on the growth of *P. noxius* cultures in this study. Ribera et al. (2016) reported that less than half of all tested *Trichoderma* isolates inhibited the radial growth rates of *P. noxius* isolates associated with landscape trees in Hong Kong. Similarly, Schwarze et al. (2012) reported that only one out of 16 possible combinations of *P. noxius* and *Trichoderma* isolates resulted in a significant decrease to the radial growth rate of the wood decay fungus compared to their respective controls. Based on similar observations, Schubert et al. (2008b) postulated that antibiosis might not be an important mechanism by which *Trichoderma* antagonizes wood decay fungi, but these results provide contrasting evidence that VOCs can significantly inhibit the growth of *P. noxius* cultures. Notably, *T. harzianum* 9132 and *T. virens* W23 caused a 68.6% and 81.3% reduction to the growth rates, respectively, of *P. noxius* KSK6 and *P. noxius* SSN3P.

The lethal effect of *Trichoderma* on *P. noxius* in dual culture tests was consistently high, except for the interaction between *P. noxius* KSK6 and *T. harzianum* 9132 (Table 5). The uniformly high lethal effect might be partially explained by the favorable habitability of MEA; other authors have reported that inhibition of wood decay fungi by *Trichoderma* in dual culture tests was significantly greater on MEA than LNA (Schubert et al., 2008b; Srinivasan et al., 1992). Specifically, Srinivasan et al. (1992) reported that soluble metabolites and volatile antibiotics were produced in significantly greater quantities by *Trichoderma* growing on MEA. As a result, the results from this test preclude discrimination among candidate *Trichoderma* isolates for biological control, but there was much greater variability in the protection afforded by various *Trichoderma* isolates to wood blocks against decay by *P. noxius*.

In this study, almost all *Trichoderma* isolates significantly inhibited wood decay by *P. noxius* compared to the controls (Table 6). Although the *Trichoderma* isolates failed to completely prevent wood decay, the sizeable reduction is in broad agreement with existing studies (Ribera et al., 2016; Schubert et al., 2008a; Schubert et al., 2008b; Schwarze et al., 2012). There are 2 possible explanations for the incomplete reduction of dry weight loss to wood blocks by *Trichoderma.* First, the wood blocks were inoculated by introducing an artificially large amount of *P. noxius* inoculum, and the disproportionate size of *P. noxius* and *Trichoderma* cultures may have favored the former. Second, a fraction of the total dry weight loss to wood blocks was probably caused by the decomposition of non-structural carbohydrates by *Trichoderma* (Klein and Eveleigh, 1998). Still, the significant reductions to the dry weight loss of wood blocks shows the strong antagonistic potential of several *Trichoderma* isolates, especially *T. harzianum* 9132 (Table 6).

After incubation for 12 weeks, the percent dry weight loss to control wood blocks inoculated with *P. noxius* KSK6 and SSN3P was much greater than reported for balsa [*Ochroma pyramidale* (Cav. ex Lam.) Urb. (Malvaceae)] wood blocks inoculated with different *P. noxius* isolates obtained from landscape trees in Hong Kong (Ribera et al., 2016), but the same values in this study are more comparable to dry weight losses reported for wood blocks removed from several landscape tree species associated with *P. noxius* in Brisbane, Australia (Schwarze et al., 2012). Collectively, these results demonstrate the importance of using species-specific wood substrates to represent the host-fungus interaction in which *Trichoderma* must intercede as a biological control agent, and this distinction gives confidence to the observations made on Senegal mahogany and rain tree wood blocks in this study.

In this study, the systematic investigation of 7 *Trichoderma* isolates demonstrated their promising antagonism towards *P. noxius*. Although the 2 *P. noxius* isolates differed in their susceptibility, several *Trichoderma* isolates showed a significant ability to inhibit growth and reduce decomposition by this aggressive wood decay fungus. None of the *Trichoderma* isolates increased the activity of *P. noxius,* and the deliberate collection of *Trichoderma* isolates from an ecological niche that resembled the desired target specificity, i.e., from basidiomycete fruiting bodies, may have been an effective selection strategy for biological control agents. In agreement with existing reports, there was no clear evidence of a primary mechanism by which *Trichoderma* antagonized *P. noxius,* and the results from different bioassays should be given equal weight in the selection of *Trichoderma* isolates. Although no single Trichoderma isolate outperformed all others in every test, there were a few that did so more than others. In particular, *T. virens* W23 produced the most chlamydospores, best inhibited the growth of *P. noxius* SSN3P, and uniquely grew faster on LNA. On the other hand, *T. harzianum* 9132 best inhibited the growth of *P. noxius* KSK6 and prevented dry weight loss to wood blocks. Still, these results are only indicative of the ability of *Trichoderma* to antagonize *P. noxius* on tree wounds, and these competitive *Trichoderma* isolates should be further examined under field conditions to confirm their efficacy. Laboratory conditions were more favorable and simplified than those governing host-fungus interaction in dynamic natural systems.

## Acknowledgments

The Agri-Food & Veterinary Authority and National Parks Board of Singapore funded this research. The contribution of laboratory assistance by Yushiella binte Ahmad, Choy May Yee, and Doris Tee Hui Ying is gratefully acknowledged.

---

**Figure 1.**
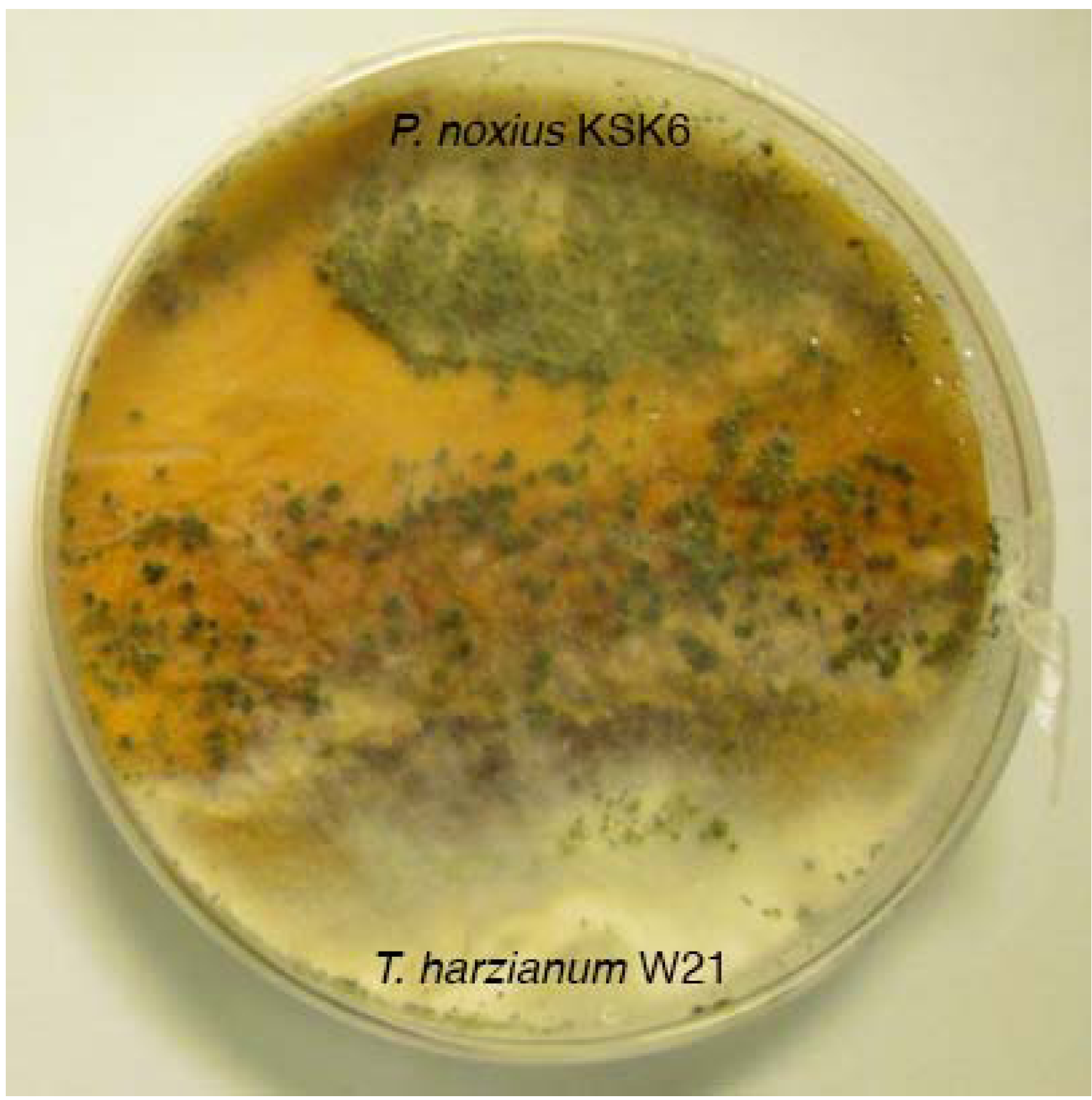
In one dual culture test, *T. harzianum* W21 (bottom) overgrew P. noxius KSK6 (top) during confrontation. The distribution of *T. harzianum* W21 over the wood decay fungus is evident by its green conidia, likely formed in response to its antagonistic interaction with P. noxius KSK6.

